# The highly abundant mRNA m^1^A modification: a new layer of gene regulation in dinoflagellates

**DOI:** 10.1101/2023.11.04.565600

**Authors:** Chongping Li, Ying Li, Jia Guo, Yuci Wang, Xiaoyan Shi, Yangyi Zhang, Nan Liang, Jie Yuan, Jiawei Xu, Hao Chen

## Abstract

The N1-methyladenosine (m^1^A) is a positively charged RNA modification known to disrupt base pairing and influence RNA stability. Despite its limited presence in the mRNA of various organism models, including yeast, mouse, and human, the exact processes of m^1^A biosynthesis, distribution, regulation, and function remain controversial. Dinoflagellates are a major group of single-celled eukaryotic phytoplankton having peculiar crystalline chromosomes. Their genes are arranged in unidirectional gene clusters along the chromosomes and only have minimal transcriptional regulation, implying the involvement of other critical regulatory mechanisms in gene expression. Here, we found that m^1^A rather than m^6^A is the most prevalent mRNA modification in dinoflagellates and asymmetrically distributed along mature transcripts. Utilizing the dinoflagellate species *Amphidinium carterae* as a study model, we identified 13481 m^1^A peaks characterized by a non-tRNA T-loop-like sequence motif within the transcripts of 10794 genes, many of which are involved in carbon and nitrogen metabolism. With enrichment around stop codon region and 3’ UTR, dinoflagellate mRNA m^1^A exhibits negative correlation with translation efficiency. Notably, nitrogen depletion (N-depletion) treatment led to significant global decrease of mRNA m^1^A amount, causing dramatic variation in translation rates with minimal changes in transcription. Additionally, our analysis uncovered distinctive methylation patterns of m^1^A modification that appears to post-transcriptionally modulate gene expression through regulating translation efficiency. Thus, our findings provide the first comprehensive m^1^A map of dinoflagellate mRNA, shedding light on its crucial role as a post-transcriptional regulatory layer to compensate the degeneration of transcriptional regulation in dinoflagellate. This study also sets the stage for further investigation into the biogenesis and functional significance of mRNA m^1^A in eukaryotes.

## Introduction

Cellular RNA undergoes over 170 covalent chemical modifications, contributing to the complexity of genetic information (Wiener & Schwartz, 2021). Despite not being the most decorated RNA species, next-generation sequencing methods facilitated the transcriptome-wide mapping of various nucleotide chemical modifications, including N6-methyladenosine (m^6^A), pseudouridine (Ψ), 5-methylcytidine (m^5^C), N1-methyladenosine (m^1^A), N4-acetylcytidine (ac^4^C), ribose methylations (Nm), and N7-methylguanosine (m^7^G) in messenger RNAs (mRNAs) (Anreiter, Mir et al., 2021, Gilbert, Bell et al., 2016, Moshitch-Moshkovitz, Dominissini et al., 2022, Roundtree, Evans et al., 2017). Of these modifications, m^6^A is the most prevalent internal mRNA modification and has garnered significant attention for its pivotal role in post-transcriptional gene regulation (Arribas-Hernández & Brodersen, 2019, Murakami & Jaffrey, 2022, Zhao, Roundtree et al., 2017).

As another form of adenosine methylation, m^1^A was initially identified in total RNA and has been extensively investigated in non-coding RNA like ribosomal RNA (rRNA) and transfer RNA (tRNA) (Dunn, 1961, Shima & Igarashi, 2020, Xiong, Li et al., 2018, Zhang & Jia, 2018). With methyl-adduct and positive charge at physiological conditions, m^1^A can block the normal base pairing with thymidine or uridine, alter local RNA secondary structure and affect the protein-RNA interactions (Xiong et al., 2018, Zhang & Jia, 2018). The mRNA m^1^A/A molar ratio ranged from 0.015% to 0.054% in mammalian cells, which was only about 10% of the level of mRNA m^6^A (Dominissini, Nachtergaele et al., 2016, Li, Xiong et al., 2016). The earlier comprehensive transcriptome-wide mapping revealed that m^1^A occurred widely on thousands of different gene transcripts, exhibited enrichment in their 5’-untranslated region (5’ UTR), changed dynamically in response to stress, and was correlated with translation enhancement (Dominissini et al., 2016, Li et al., 2016). However, subsequent studies demonstrated that cytosolic mRNAs were sparsely modified with m^1^A at very low stoichiometries, which usually contained tRNA T-loop like structures, led to translation repression and were invariably catalyzed by TRMT6/TRMT61A complex (Safra, Sas-Chen et al., 2017, Schwartz, 2018). Another study also reported that m^1^A was a rare internal mRNA modification and only appeared in one mitochondrial transcript (Grozhik, Olarerin-George et al., 2019). Thus, due to the limited number of gene transcripts bearing m^1^A sites, the biological function of these internal m^1^A modifications in mammalian cells awaits further verification. In *Petunia hybrida,* a plant species containing relatively high amounts of m^1^A (0.25-1.75% of total adenines in mRNA), m^1^A modifications were identified in 3231 genes, mainly located in coding sequence near start coding regions, and showed dynamic distribution upon ethylene treatment (Yang, Meng et al., 2020). However, the role of these m^1^A modifications in post-transcriptional gene regulation remains largely unexplored.

Dinoflagellates, consisting of important marine primary producers, harmful bloom-forming microalgae and photo-symbionts of marine invertebrates, are a major group of unicellular eukaryotic marine phytoplankton. Their unique characteristics, including permanently condensed liquid-crystalline chromosomes lacking nucleosomes (Wisecaver & Hackett, 2011, Wong, 2019), alternating unidirectional gene arrays (Nand, Zhan et al., 2021, Nelson, Hazzouri et al., 2021, Shoguchi, Shinzato et al., 2013, Stephens, González-Pech et al., 2020), mRNA maturation with trans-splicing (Lidie & Van Dolah, 2007, Zhang, Hou et al., 2007), and non-typical promoter harbouring no TATA box (Guillebault, Sasorith et al., 2002, Lin, Cheng et al., 2015), distinguish them from other eukaryotes in terms of chromatin organization and gene expression. Instead of controlling gene expression at transcriptional level, it was believed that dinoflagellates mainly rely on post-transcriptional regulation (Roy & Morse, 2013, Zaheri & Morse, 2022). There is a clear paucity of both basal and promoter-specific transcription factors in dinoflagellates genomes, which encode only half number of expected components of RNA polymerase II (RNAP II) and less than 10% of gene specific transcription factors compared to classical eukaryotes (Roy & Morse, 2013, Zaheri & Morse, 2022). Numerous studies also indicated that for different dinoflagellate species including *Symbiodinium* sp., *Lingulodinium* sp., *Scrippsiella* sp., *Alexandrium* sp., only a few genes exhibited significant changes in their transcript abundance under light-dark transition, temperature change, circadian rhythm and stressful conditions (Chen, Dougan et al., 2023, Moustafa, Evans et al., 2010, Roy, Beauchemin et al., 2014, Yang, Beszteri et al., 2011, Zaheri, Dagenais-Bellefeuille et al., 2019).

As stated above, there is still controversy regarding the existence and role of m^1^A in mRNA. Dinoflagellates lack transcription control over gene expression and primarily depend on post-transcriptional regulation. In this study, we found unexpectedly high m^1^A level in mRNA of dinoflagellates, making them an ideal model system to investigate the function of m^1^A within transcripts. We identified thousands of m^1^A-modified transcripts present in dinoflagellate transcriptome, which showed marked difference with typical eukaryotes in terms of the major near stop codon and 3’ UTR distribution pattern, and non-T loop-structure like motif. m^1^A methylation seemed to regulate mRNA fates through decreasing translation efficiency and may be utilized to cope with nutrient stress. These findings clearly demonstrated that m^1^A governs mRNA translation rates and adds another layer of gene expression control at post-transcriptional level in dinoflagellates.

## Results

### Dinoflagellate mRNA is highly decorated by m^1^A modification

Due to the cholesteric liquid crystalline structure of chromosomes, extensive use of 5’-trans-splcing in mRNA and reduced role of transcriptional regulation in dinoflagellates (Fig. 1A), we envisaged that RNA modification may be involved in post-transcriptional regulation of their gene expression. Therefore, we first investigated the frequently occurred RNA modifications in dinoflagellate *A. carterae* and the results showed that all RNA modifications including m^1^A, m^6^A, m^6^Am and m^7^G were readily detected in total cellular RNA, while only m^6^A and m^1^A could be detected with considerable level in their poly(A)+ mRNA fractions (Supplementary Fig. 1). Moreover, the dot blotting analysis using the m^1^A-specific antibody also demonstrated that *A. carterae* contains much more m^1^A in their mRNA than mammalian HEK293T cell lines (Fig. 1B).

**Figure 1.**
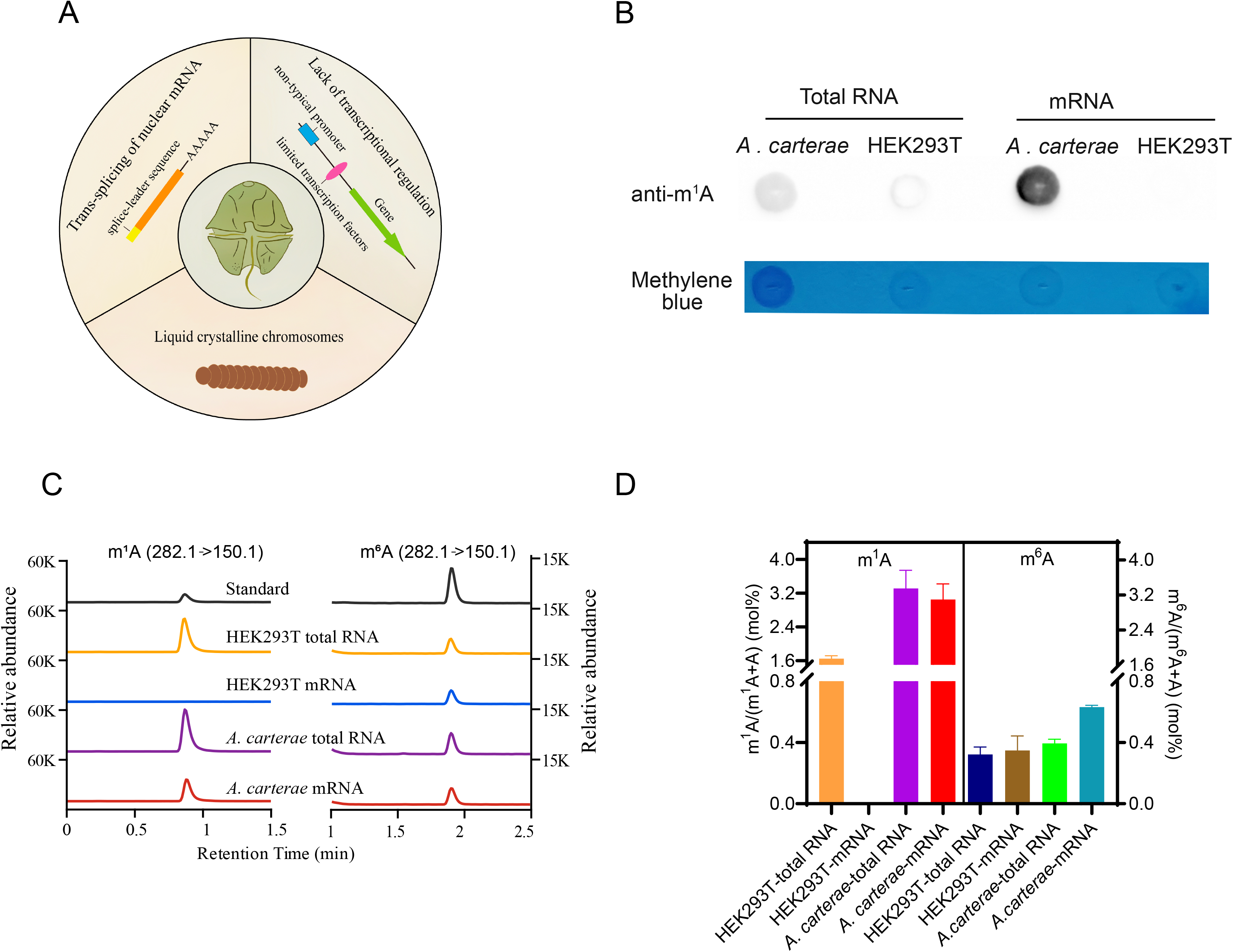
Quantitative detection of internal m^1^A sites in dinoflagellate *A. carterae*. (A) Several unusual features in dinoflagellates; (B) and (C) Dot-blot assay of m^1^A abundance in both total RNA and mRNA of *A. carterae* and HEK 293T cell lines respectively. (D) LC-MS/MS chromatograms and quantification of m^1^A and m^6^A levels in the mRNA and total RNA isolated from *A. carterae* and HEK 293T cell lines. Mean values ± SE are shown; n =3.

Consistent with the very low amount of m^1^A in mammalian cells in previous studies (Dominissini et al., 2016, Li et al., 2016), liquid chromatography–tandem mass spectrometry (LC-MS/MS) based quantification showed that m^1^A level is barely detected (near to baseline) in HEK293T mRNA, while m^1^A accounts for about 1.65% of the total adenines in its total RNA (Fig. 1C and 1D). Surprisingly, the m^1^A content in *A. carterae* mRNA reached up to 3.05% of the total adenines (Fig. 1C and 1D), which is close to the m^1^A level of its total RNA and two orders of magnitude higher than reported amounts of m^1^A in mammalian cells (Dominissini et al., 2016, Li et al., 2016). Our results also revealed that the m^6^A constitutes about 0.63% of the total adenine in *A. carterae* mRNA (Fig. 1C and 1D), which is only equivalent to one fifth of the corresponding m^1^A level and slightly higher than m^6^A amount of HEK293T cells (∼0.35%). The relatively higher amount of m^1^A than m^6^A level in their mRNA was also observed in other dinoflagellates including *Crypthecodinium cohnii* and *Symbiodinium* sp., while not observed in unicellular green algae *Chlamydomonas reinhardtii* (Supplementary Fig. 2). Thus, this proportion of mRNA m^1^A in dinoflagellates is likely conserved among different taxa of dinoflagellates and is much higher than other eukaryotes. Furthermore, it was uncovered that abundant m^6^A resides in poly(A) tails of *Trypanosoma brucei* mRNA (Viegas, de Macedo et al., 2022), we asked whether dinoflagellate m^1^A modifications are also located within mRNA poly(A) tails. The results showed that these m^1^A sites are mainly distributed in the non-poly(A) region instead of poly(A) tails of the mature transcripts (Supplementary Fig. 3).

### MeRIP-Seq unveils m^1^A profile throughout the transcriptome of Dinoflagellate *A. carterae*

To gain further insight into the roles of m^1^A, we sought to characterize m^1^A sites at a transcriptome-wide level by using anti-m^1^A antibody-based MeRIP-seq (m^1^A-specific methylated RNA immunoprecipitation). A total of 13481 peaks in the mRNAs of 10794 genes were detected in the *A. carterae* under normal growth condition (Fig. 2A). Each methylated gene exhibits an average of 1.25 peaks, and 80.66%, 15.21% and 3.19% of these transcripts contain one peak, two peaks and three peaks, respectively (Fig. 2A). Only a few transcripts contain four or more peaks (Fig. 2A). Further analysis showed that the majority of these peaks are strongly enriched in the stop codon segment and 3’ UTR with two peaks positioned near the stop codon (Fig. 2B, 2C and 2D). Specifically, 39.2% and 37.2% of these m^1^A peaks are distributed in stop codon segment and 3’ UTR of the encoded transcripts, while only 20.1%, 1.9% and 1.6% of these peaks are located in CDS, 5’ UTR and Start Codon segment respectively (Fig. 2B).

**Figure 2.**
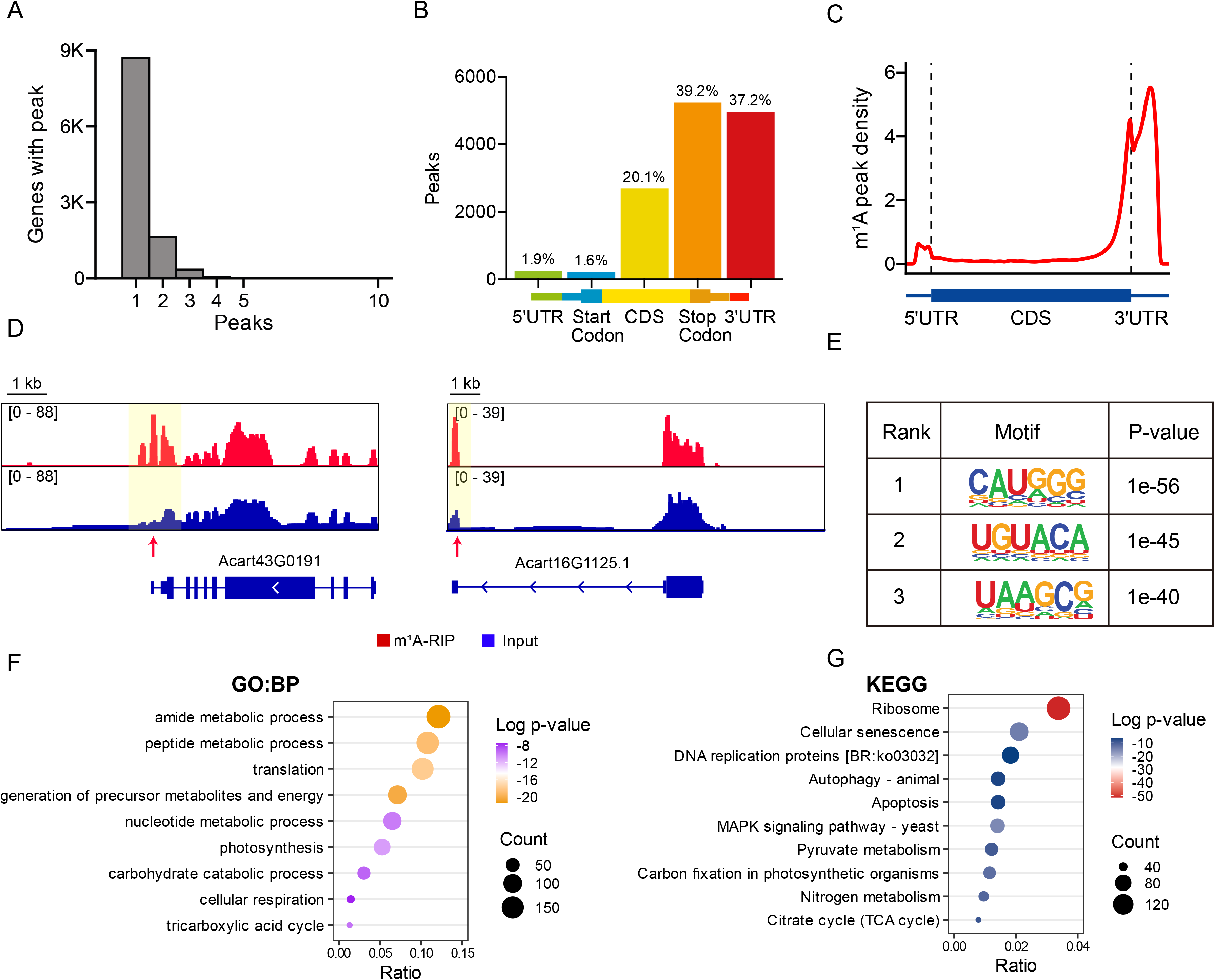
The transcriptome-wide landscape of m^1^A methylome in *A. carterae*. (A) The detailed gene numbers of m^1^A-methylated transcripts exhibiting 1, 2, 3, or 4+ peaks. (B) Percentage of m^1^A peaks in different regions of the mRNA. (C) Metagene profile of m^1^A peaks along the full-length transcripts. The 5L UTR, CDS and 3L UTR were normalized to their average length according to the *A. carterae* genome annotation files. (D) Examples of m^1^A modified genes in *A carterae*. (E) The potential motifs for m^1^A-containing peak regions. (F) and (G) GO and KEGG analyses of m^1^A-methylated genes.

Clustering of all enriched m^1^A peak sequences revealed that these peaks in dinoflagellate *A. carterae* shared common sequence motifs including CAUGGG (E value = 1e-56), GCCACGC (E value = 1e-45) and UAAGCG (E value = 1e-40) (Fig. 2E), which are apparently different from previous tRNA T-loop-like elements GUUCNANNC of m^1^A peaks found in mammalian cells (Li, Xiong et al., 2017, Safra et al., 2017). Furthermore, we cloned the *A. carterae* homologs of putative tRNA methyltransferase and purified the recombinant protein complex of AcTRMT6/AcTRMT61A (Supplementary Fig. 4A). The *in vitro* biochemical assay indicated that AcTRMT6/AcTRMT61A cannot catalyze the production of m^1^A in both *A. carterae* and HEK293T mRNA (Supplementary Fig. 4B). In contrast, new m^1^A synthesis was observed in total RNA of these cells and *A. carterae* small RNA (Supplementary Fig. 4B). These results demonstrated that dinoflagellate mRNA m^1^A methylation is installed by a yet unknown methyltranferase, which is distinct from typical m^1^A methyltransferase of eukaryotic tRNAs.

To investigate the biological functions of m^1^A-methylated mRNAs, we next performed GO and KEGG enrichment analysis. GO analysis indicated that the top ranked lists of enriched GO categories of these m^1^A-decorated transcripts are correlated with amide biosynthetic process, peptide metabolic process, translation, generation of precursor metabolites and energy, nucleotide metabolic process, photosynthesis etc. (Fig. 2F). Meanwhile, KEGG enrichment analysis also revealed the association between m^1^A -modified genes and similar pathways such as ribosome, carbon fixation in photosynthetic organisms, nitrogen metabolism, pyruvate metabolism and others (Fig. 2G). These results implied an important role of m^1^A in translation regulation and cellular carbon/nitrogen metabolism.

### m^1^A associates with translation inhibition

Bearing in mind little transcriptional regulation in dinoflagellates and interference effects of m^1^A methylation on base pairing and RNA-protein interaction, we then assessed possible function of m^1^A in post-transcriptional regulation within these organisms. Our analysis showed that m^1^A-methylated genes in dinoflagellate *A. carterae* have significantly higher mRNA levels than the unmethylated ones (Fig. 3A). Particularly, except for the 5’ UTR with m^1^A peaks, the transcript abundance is positively correlated with the presence of m^1^A peaks in other gene regions (Fig. 3B). Short poly(A) tails length has been recently regarded as a conserved feature of the highly expressed genes across eukaryotes (Lima, Chipman et al., 2017, Passmore & Coller, 2022). Consistent with this notion, we also found that the length of mRNA poly(A) tails is negatively correlated with mRNA abundance (Supplementary Fig. 5A) and m^1^A-marked transcripts possessed shorter poly-A tails in *A. carterae* (Fig. 3C). Moreover, the mRNAs decorated with m^1^A tend to have lower GC content when compared with non-methylated mRNAs (Fig. 3D). However, with the exception of 5’ UTR, GC content reduction is only observed in both CDS and 3’ UTR of m^1^A-modified transcripts (Supplementary Fig. 5B). Accordingly, the structural potential predicted by ViennaRNA in near stop codon region with or without m^1^A modification showed that m^1^A-methylated regions displayed a lower probability to form stable secondary structures than nonmethylated counterpart (Fig. 3E). These results suggested that m^1^A preferred to install on highly expressed genes and their unstructured mRNA regions.

**Figure 3.**
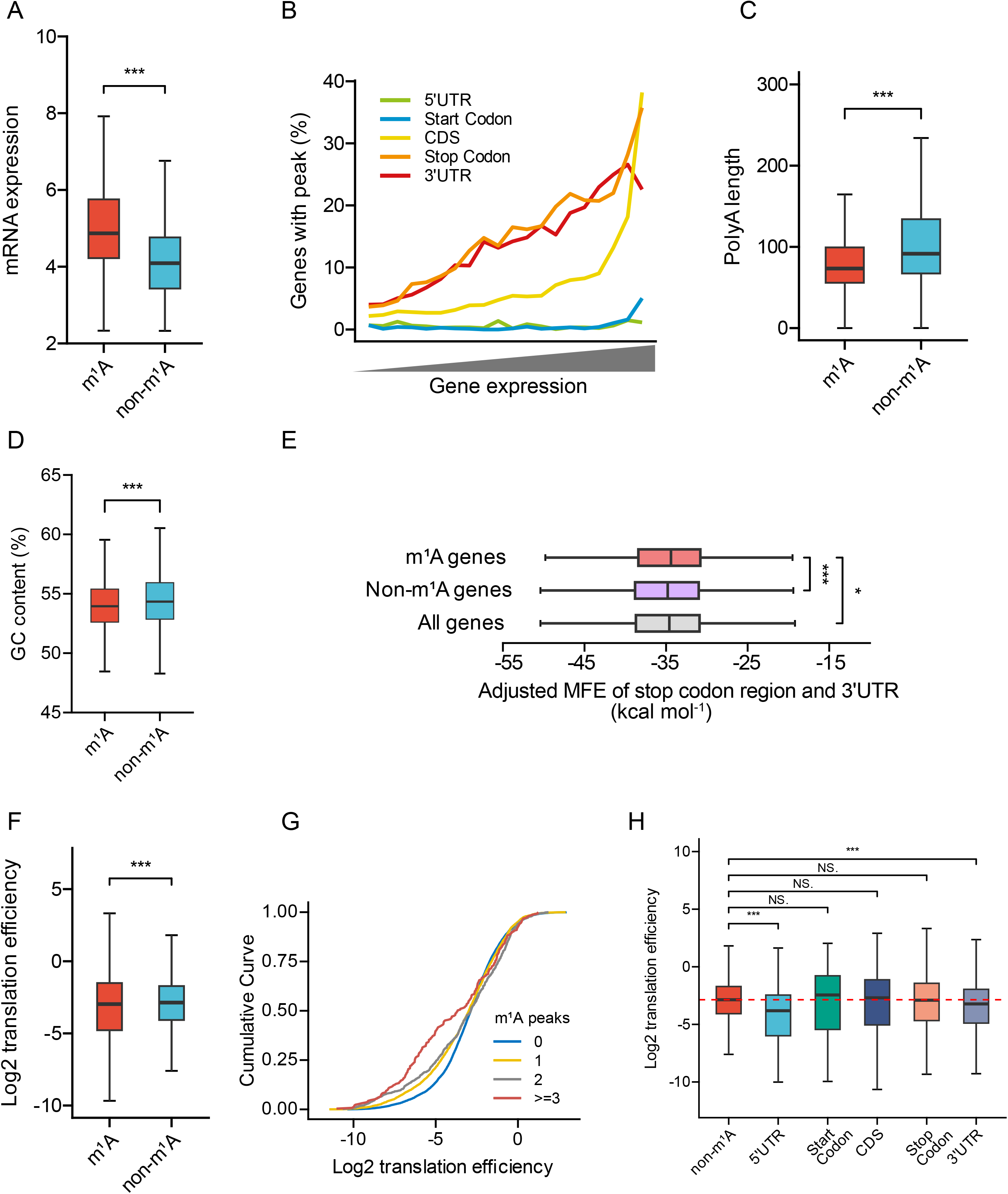
m^1^A associates with mRNA abundance, gene structure and regulates translation efficiency. (A) and (B) The correlation of m^1^A modifications with gene expression level. (B) Gene expression level plots against different regional methylation. (C-F) A comparison between the median poly(A) length (C), GC content (D), minimum folding energy (MFE) (E), and translation efficiency (F) of genes with or without m^1^A modifications. (G) Cumulative distribution of Log2-fold changes of translation efficiency of the m^1^A-modified genes with different amount of m^1^A peaks. (H) With same data from (F), translation efficiency was further compared among non-m^1^A-methylated genes with m^1^A-modified genes occurred in different mRNA segments. For all box plots, the median values in each group are marked by a center line. The boxes upper and bottom boundaries denote the first quartile and third quartile, while the error bars show the highest and lowest values. p values were determined using Wilcox test. ns, not significant; *p < 0.05 and ***p < 0.001.

We further explored whether m^1^A methylation would affect the decoding process of endogenous transcripts. Comparative analysis of translation efficiency for transcripts with or without m^1^A indicated that m^1^A-modified transcripts are associated with decreased translation efficiency (Fig. 3F). Consistently, the transcripts with more m^1^A peaks exhibit much lower translation efficiency (Fig. 3G). Interestingly, this significant translation inhibition is only occurred in mRNAs with methylated 5’ UTR or 3’ UTR regions (Fig. 3H, Supplementary Fig. 5C and 5D). Thus, our results implied that the inhibitory effect of m^1^A modification on transcript translation is context-dependent.

### Dynamic m^1^A methylation across stress condition

Given the involvement of m^1^A methylated genes in nitrogen metabolism, we further explored how m^1^A methylation level and distribution responds to N-depletion stress. Consistent with previous studies (Lai, Yu et al., 2011, Li, Li et al., 2021), the growth of *A. carterae* cells was also severely inhibited under N-depletion condition (Fig. 4A). RNA-seq analysis results indicated that there were only a total of 134 differentially expressed genes (DEGs) upon N-deficiency, of which 81 and 53 exhibited significant up-regulation and down-regulation respectively (Supplementary Fig. 6A), which further substantiate the lack of gene expression control at transcriptional level. The up-regulated DEGs were mainly engaged in GO terms such as pigment biosynthetic process, organic cyclic compound biosynthetic process, nitrate assimilation and photosynthesis, and also KEGG pathways related to photosynthesis, nitrogen metabolism and pyruvate metabolism (Supplementary Fig. 6B and 6C). The down-regulated DEGs are involved in GO terms associated with phosphate ion transport and secondary metabolic process, and KEGG pathways including ABC transporters, cholesterol metabolism and also nitrogen metabolism (Supplementary Fig. 6B and 6C).

**Figure 4.**
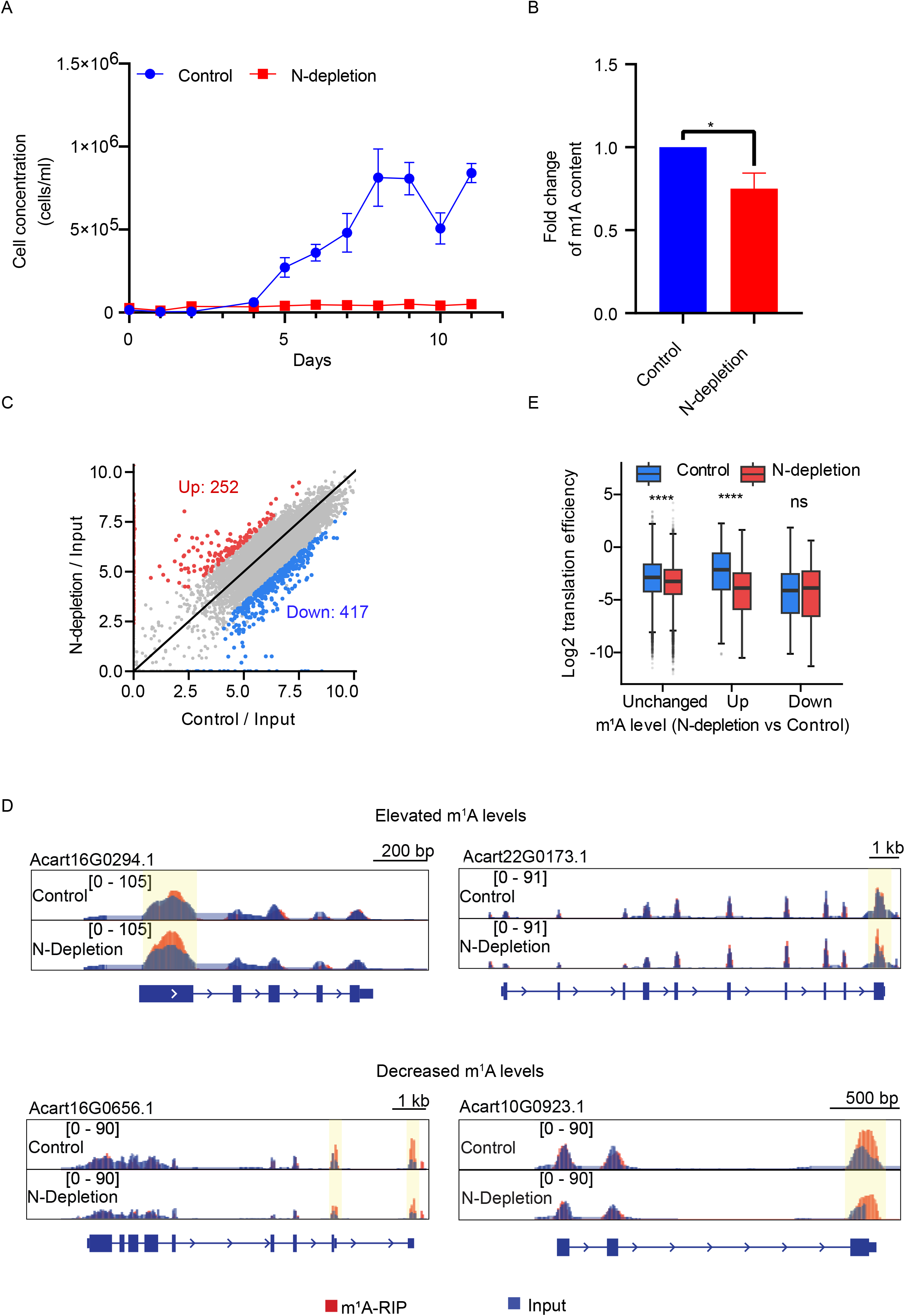
Characterize the role of m^1^A upon nitrogen deficiency. (A) Growth curves of *A. carterae* cells under different nitrogen conditions. (B) The fold change of mRNA m^1^A level of *A. carterae* cells grown under nitrogen-depletion conditions for 6 days (with paired t-test, p* < 0.05). All the data are means ± SE, n =3. (C) Comparison scatter plot of MeRIP analyses of *A. carterae* cells in N-depleted vs N-replete treatments. Red dots denote stress-upregulated peaks; blue dots denote N-depletion downregulated peaks. (D) Representative examples of elevated and reduced m^1^A peaks upon N-depletion. (E) Box plot shows the translation efficiency changes of genes with or without differential methylation identified in (C). Boxes represent 25th–75th percentile (line at the median) with whiskers at 1.5 × interquartile range (Wilcox test, ****p < 0.0001).

Recent studies suggested mRNA translation rate change in dinoflagellate *Lingulodinium polyedra* was an important post-transcriptional mechanism for circadian rhythm adaptation (Bowazolo, Song et al., 2022). Inspired by these findings, we then utilized Ribo-seq to determine whether the change of mRNA translation efficiency was used to cope with N-deficiency. From the Ribo-seq data, we found that 1313 and 2474 genes showed significantly elevated and reduced translation efficiency respectively (Supplementary Fig. 6D). The genes with increased translation efficiency are mainly responsible for regulating autophagy, ABC transporters, lysosome, amino sugar and nucleotide sugar metabolism (Supplementary Fig. 6E and 6F), while the genes with decreased translation efficiency are involved in affecting purine ribonucleotide metabolic process, inorganic ion transmembrane transport, nucleotide metabolic process, photosynthesis, carbohydrate catabolic process, small molecule biosynthetic process, and translation factors (Supplementary Fig. 6E and 6F). With much more change in translation efficiency than gene expression level, our data demonstrated that dinoflagellate adapt to N-depletion condition mainly through governing gene translation rates.

Next, we measured the m^1^A content of *A. carterae* cells grown in N-limitation condition and found that their global mRNA m^1^A level significantly decreased to 75% of that in the normal growth condition (Fig. 4B). MeRIP-seq analysis were further performed to reveal the potential role of m^1^A modification for dealing with nutrient stress. 10569 high-confidence m^1^A peaks were observed in 8853 genes under N-depletion condition. Differential RNA methylation analysis of the MeRIP-seq data indicated that 252 and 417 genes display significantly increased and decreased m^1^A methylation respectively upon N-deficiency (Fig. 4C and 4D), of which only 21 genes belonged to the DEGs at the transcriptional level (Supplementary Fig. 7A). Specifically, the increased m^1^A sites were mainly distributed on both CDS and 3’ UTR, while reduced m^1^A level primarily occurred on Stop codon region and 3’ UTR segment (Supplementary Fig. 7B). In order to clarify the function of differentially methylated transcripts, GO and KEGG enrichment analyses were further conducted. With no functional enrichment of the genes with decreased m^1^A methylation, the genes with enhanced m^1^A methylation showed regulation on translation, amide biosynthetic process and nitrogen metabolism (Supplementary Fig. 7C and 7D). These results suggested that m^1^A modification was also utilized for post-transcriptional regulation of translation and nitrogen metabolism in dinoflagellates.

Translation efficiency of the differentially methylated mRNA was further compared between normal and N-depletion condition. In consistent with the translation repression effect of m^1^A modification, m^1^A transcripts with diminished methylation have relatively higher translation efficiency, and augmented m^1^A methylation levels in transcripts lead to translation inhibition (Fig. 4E, Supplementary Fig. 7E, 7F and 7G). Thus, the above results implied that m^1^A mediated-gene expression control via translation efficiency regulation might be a critical strategy for dinoflagellate adaption under N-depletion condition.

## Discussion

N1-methylation of RNA adenosine (m^1^A) normally occurs at a much lower abundance than the most plentiful m^6^A modification in cellular mRNA of typical eukaryotes (Dominissini et al., 2016, Li et al., 2016, Wiener & Schwartz, 2021). Intriguingly, in contrast to the 5’ UTR enrichment pattern and low abundance reported in early studies, we found m^1^A is predominant in dinoflagellates mRNA and mostly installed within stop codon region and 3’ UTR (Supplementary Fig. 8), which resembles the m^6^A peaks located across the CDS-3’ UTR junctions along the transcript length in typical eukaryotes (Dominissini, Moshitch-Moshkovitz et al., 2012, Meyer, Saletore et al., 2012). Notably, the involvement of m^1^A-methylated transcripts in cellular translation, carbon and nitrogen metabolism indicates that m^1^A-meidated post-transcriptional regulation represents an important layer of controlling critical biological processes in dinoflagellates. m^6^A modification is added at nascent pre-mRNA stage, and then preserved until mRNA maturation (Ke, Pandya-Jones et al., 2017). It remains to be investigated when and how these m^1^A are deposited in mRNA within cells (Supplementary Fig. 8). Given the similar sequence element with tRNA T-loop, tRNA m^1^A methyltransferase complex TRMT6/61A is also responsible for the generation of the scarce cytosolic mRNA m^1^A modification in mammalian cells (Li et al., 2017, Safra et al., 2017). Instead, the inability to produce m^1^A methylation in *A. carterae* mRNA by the TRMT6/61A heterocomplex suggests that there may be a new kind of methyltransferase in dinoflagellates.

As an isomer of m^1^A, m^6^A can stabilize RNA local structure by strengthening base stacking (Roost, Lynch et al., 2015). In comparison, the methyl group of m^1^A modification is located at the Watson-Crick base-pairing edge, which results in base-pairing block and local duplex melting (Helm, Giegé et al., 1999, Zhou, Kimsey et al., 2016). Generally, mRNA harboring m^6^A sites have short half-lives and decreased stability (Wang, Lu et al., 2014, Zaccara & Jaffrey, 2020). Given the limited global transcriptional change of dinoflagellate genes, the contribution of m^1^A to their mRNA stability maintenance is probably neglectable. m^6^A-dependent RNA structural remodeling was known to affect the mRNA abundance as well as alternative splicing through modulating RNA-protein interactions (Liu, Dai et al., 2015). When localized in the CDS, m^6^A positively promotes translation rates by hindering the formation of stable secondary structures (Mao, Dong et al., 2019). Besides, 5’ UTR m^6^A is also able to drive cap-independent translation in the absence of 5’ cap-binding proteins (Meyer, Patil et al., 2015). However, there is no agreement regarding the effect of m^1^A on translation efficiency (Dominissini et al., 2016, Safra et al., 2017). Here, we demonstrated that dinoflagellate m^1^A preferentially occurs on highly expressed genes and was negatively associated with translation rates of transcripts. Unlike other regions, methylation at either 5’ UTR or 3’ UTR showed a negative correlation with translation efficiency, suggesting that regional effect of m^1^A modification might also exist within dinoflagellate mRNA.

microRNA has been reported to show differential expression in dinoflagellates under stress conditions, suggesting it could be used to regulate gene expression at the post-transcriptional level (Shi, Lin et al., 2017). Recent study on m^1^A profiling in primary neurons has shown that m^1^A modification in the 3’ UTRs of mRNA could prevent the binding of miRNA, resulting in enhanced gene expression (Baumgarten, Bayer et al., 2013, Zhang, Yi et al., 2023). microRNA generally mediates transcript degradation and/or translation repression by binding to 3’ UTR of mRNA (Shang, Lee et al., 2023, Song, Li et al., 2019). Given the high stability of dinoflagellate mRNA, which has a median half-life of 33 hours (Morey & Van Dolah, 2013), it seems unlikely that microRNA-dependent post-transcriptional regulation also accounts for the m^1^A-mediated translation inhibition. Many regulatory sequences involved in translation initiation control are embedded in 3’ UTR (Andreassi & Riccio, 2009, Mayr, 2019). The m^6^A reader proteins YTHDF1/2 binds m^1^A-methylated transcripts with comparable affinity to m^6^A-recognition activity (Seo & Kleiner, 2020). Another possibility is that m^1^A in 3’ UTR or m^1^A reader protein(s) interacts with translation initiation factor to destabilize closed-loop formation and then stall translation.

Given the paucity of transcription factors and differentially expressed genes at transcriptional level, dinoflagellates preferentially control gene expression through post-transcriptional regulation (Roy & Morse, 2013, Zaheri & Morse, 2022). Despite lacking significant difference in the transcriptome over a daily light-dark cycle, dinoflagellates *Lingulodinium* and *Symbiodinium* possess coordinated changes in the number of ribosome-protected fragments (Bowazolo & Morse, 2023, Bowazolo et al., 2022), suggesting translational control serves as a major strategy for regulating gene expression. However, the control mechanism behind translation rates change remained to be addressed. In current study, our results also revealed that *A. carterae* mainly responded to N-depletion by altering mRNA translation rates (Supplementary Fig. 8). Moreover, we found that m^1^A could act as a delicate brake to reduce the translation efficiency of these transcripts relating with metabolism and protein synthesis. Therefore, dynamic m^1^A methylations in mRNA enables dinoflagellates to fine-tune the regulatory strength of translation machinery in response to various environmental stimuli. It should be noted that there is also similar level of m^6^A like normal eukaryotes within dinoflagellate mRNA. The division of labor between these two marks and cross-talk with other cellular pathways in mRNA post-transcriptional regulation remains to be further investigated.

## Acknowledgments

We would like to thank Ms. Hui Zhou for her help in recombinant protein purification. The authors would also like to acknowledge the technical support from Hua Li and Lin Lin at SUSTech CRFT. This work was supported by Center for Computational Science and Engineering at Southern University of Science and Technology.

This work was supported by National Key Research and Development Program of China (2022YFC2702705), National Natural Science Foundation of China (32170604 to H.C.). This work was also supported by Pearl River Recruitment Program of Talents (2021QN02Y122) and Department of Health of Guangdong Province (B2021032) to H.C., Shenzhen Key Laboratory of Gene Regulation and Systems Biology (Grant No. ZDSYS20200811144002008) from Shenzhen Innovation Committee of Science and Technology and Funding for Scientific Research and Innovation Team of The First Affiliated Hospital of Zhengzhou University (ZYCXTD2023004).

## Author contributions

C.P.L., J.W.X and H.C. designed and conceived the experiments. C.P.L., Y.L., J.G., and N.L. performed most of the experiments. Under the supervision of J.W.X. and J.Y., Y.C.W., Y.L. and J.G. analyzed the NGS data. X.Y.S. and Y.Y.Z. assisted in LC-MS/MS experiments. C.P.L. and H.C. wrote the manuscript with the inputs from other authors. All authors have read and approved the final manuscript.

## Supplementary figure legends

**Figure S1.**
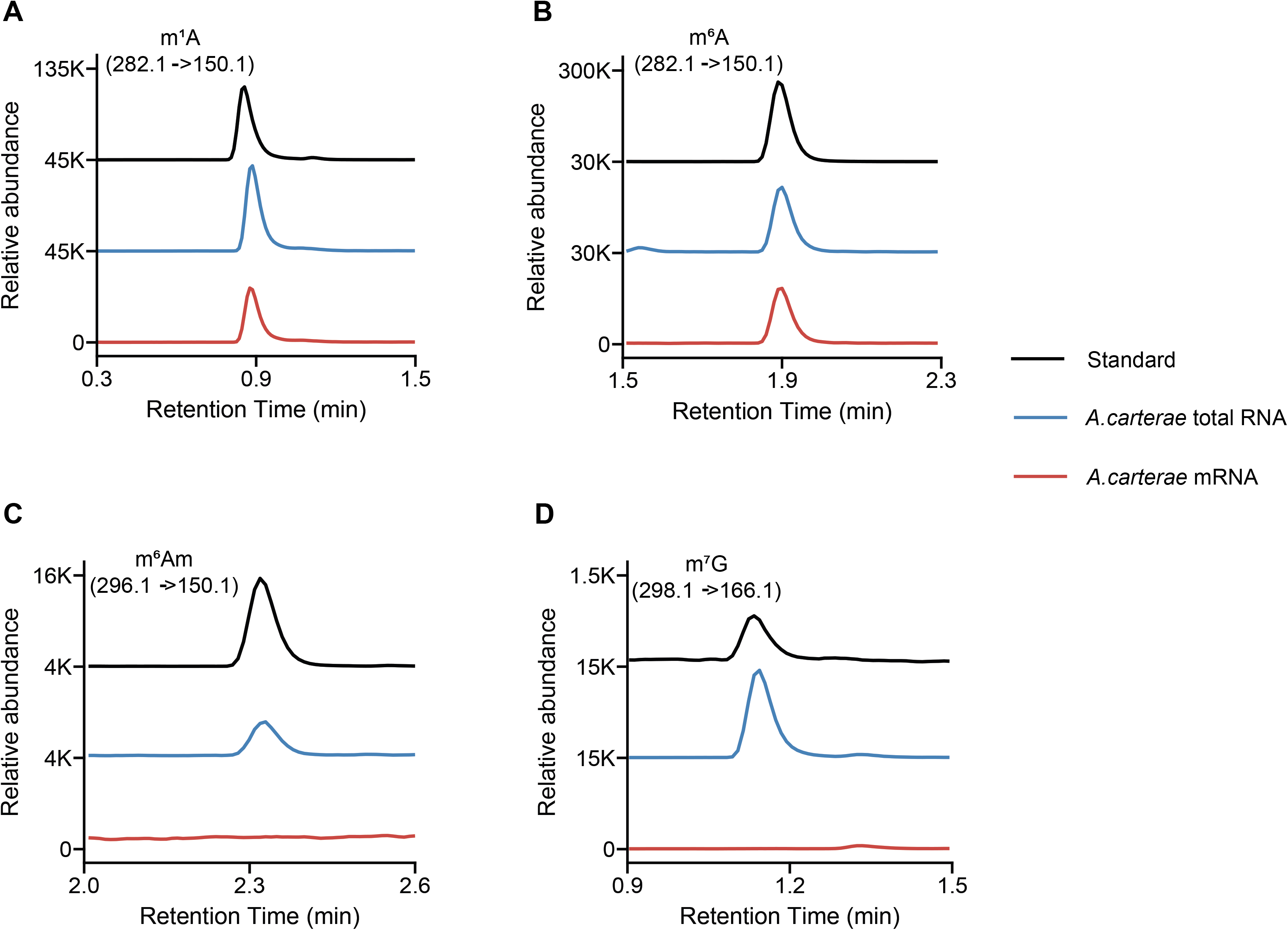
Common RNA modification species detected in isolated total RNA and mRNA of dinoflagellate *A. carterae*. (A-D) RNA spectra of m^1^A, m^6^A, m^6^A_m_ and m^7^G present in standard samples, *A. carterae* total RNA and mRNA respectively.

**Figure S2.**
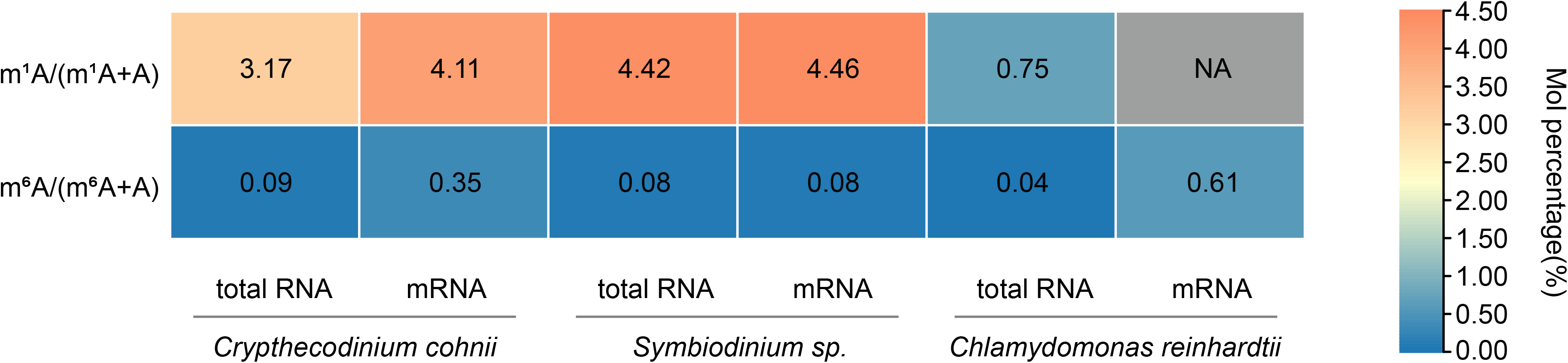
Quantitative analysis of m^1^A and m^6^A level in total RNA and mRNA in other dinoflagellates and green algae *Chlamydomonas reinhardtii*. NA, abbreviation of ‘not available’, means the m^1^A level in mRNA of *Chlamydomonas reinhardtii* is below the detection limitation of standards. Sample size = 2 biological replicates for each group.

**Figure S3.**
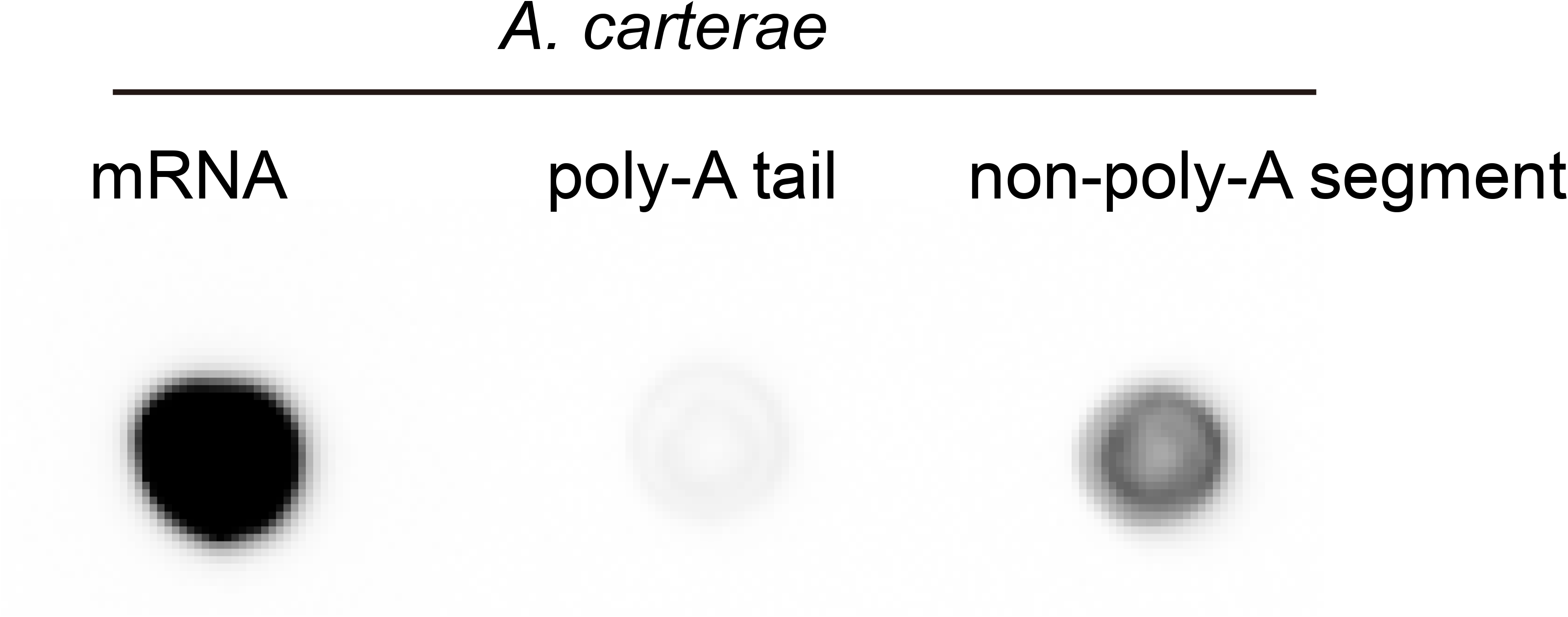
Distribution of m^1^A in mRNA poly(A) tails and non-poly(A)-segments

**Figure S4.**
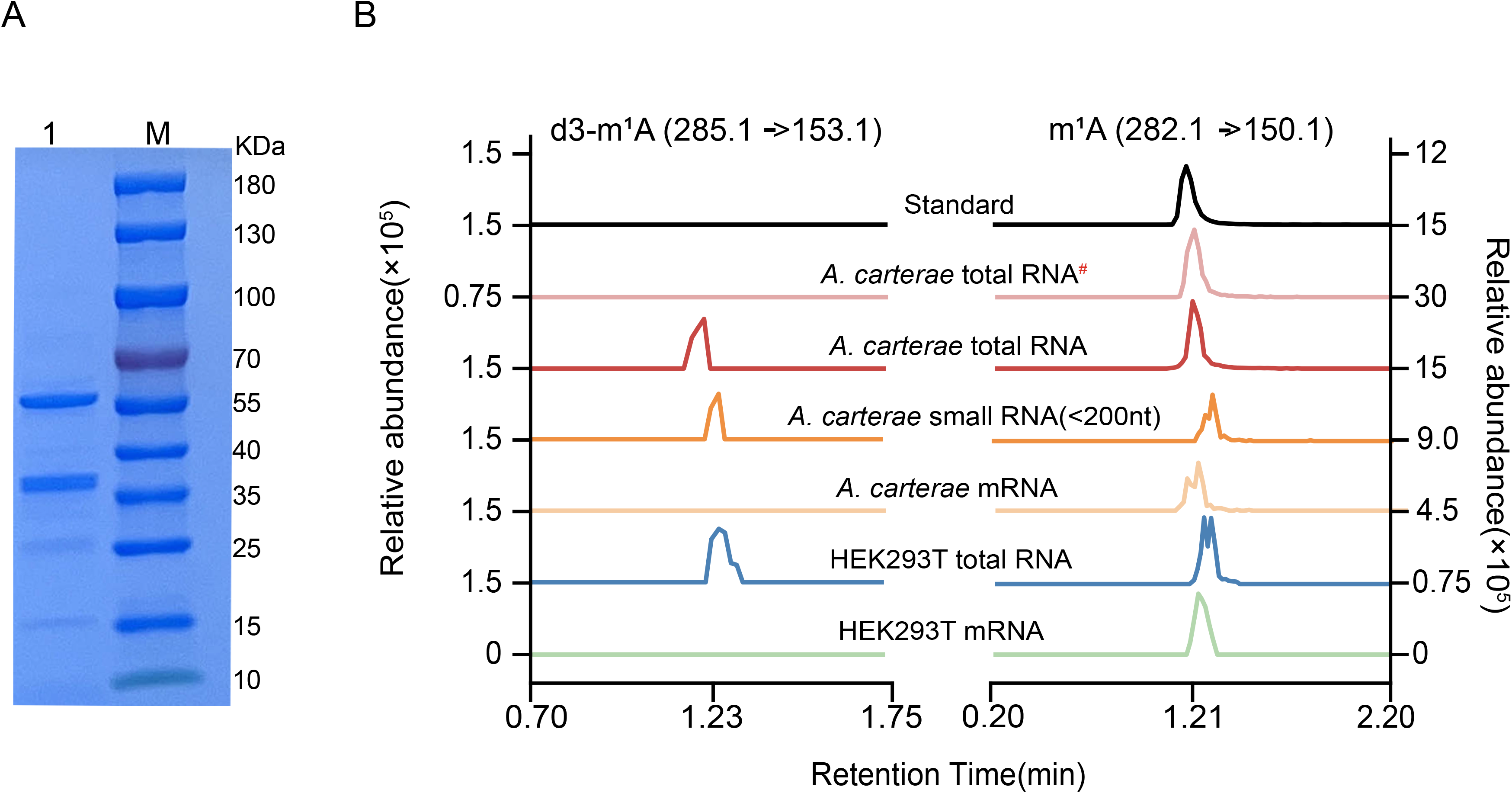
The methyltransferase activities of AcTRMT6/AcTRMT61A heterocomplex to produce new m^1^A sites using different RNA substrates. (A) SDS-PAGE analysis showing the purified AcTRMT6/AcTRMT61 complex with expected molecular weights of 38.5 kDa and 51.5 kDa respectively. Lane 1, the purified recombinant AcTRMT6/AcTRMT61A heterocomplex; Lane M, the Thermo Scientific PageRuler Plus Prestained Protein Ladder. (B) LC-MS/MS spectra of various RNA substrates from in vitro methylation reaction. #, represents the addition of heat-inactivated AcTRMT6/AcTRMT61A heterocomplex in this reaction mixture. The native recombinant protein complex AcTRMT6/AcTRMT61A were used in all other reactions.

**Figure S5.**
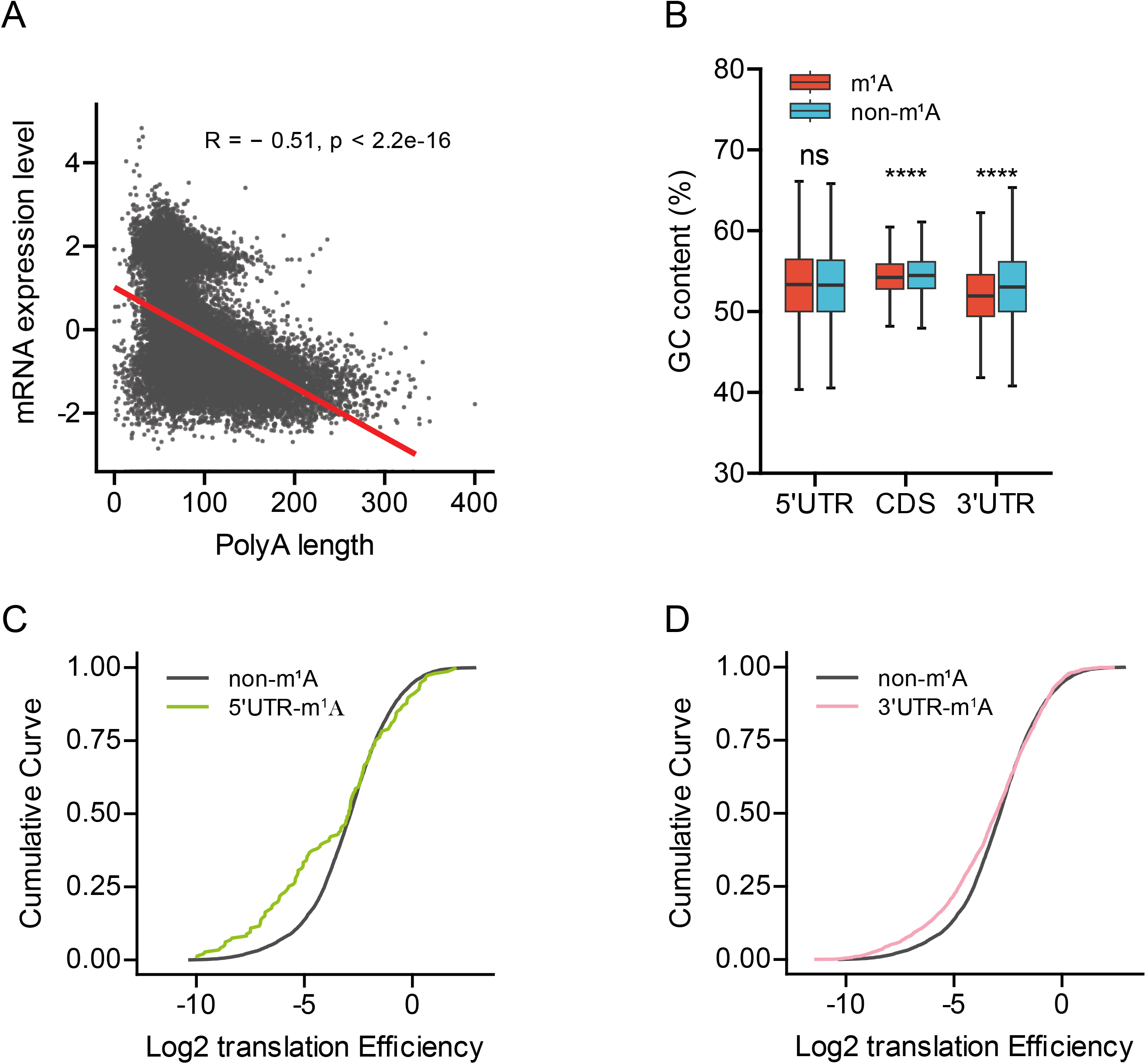
The correlation between gene expression level and poly(A) tails, and detailed characteristics of m^1^A-methylated transcripts. (A) Correlation analysis of mRNA abundance with the poly(A) length of encoded transcripts in *A. carterae*. (B) A comparison of GC contents of 5’ UTR, CDS and 3’ UTR of m^1^A-methylated genes with the counterparts of unmethylated genes (Wilcox test, ns = not significant; *p <0.05; ****p < 0.0001). Median of GC content in each group is indicated by a center line, the box shows the upper and lower quantiles, whiskers show the 1.5× interquartile range. The outliers are not shown. (C) and (D), The Log2 translation efficiency is plotted as accumulative fractions for transcripts of both non-methylated and 5’ UTR-methylated or 3’ UTR-methylated genes respectively.

**Figure S6.**
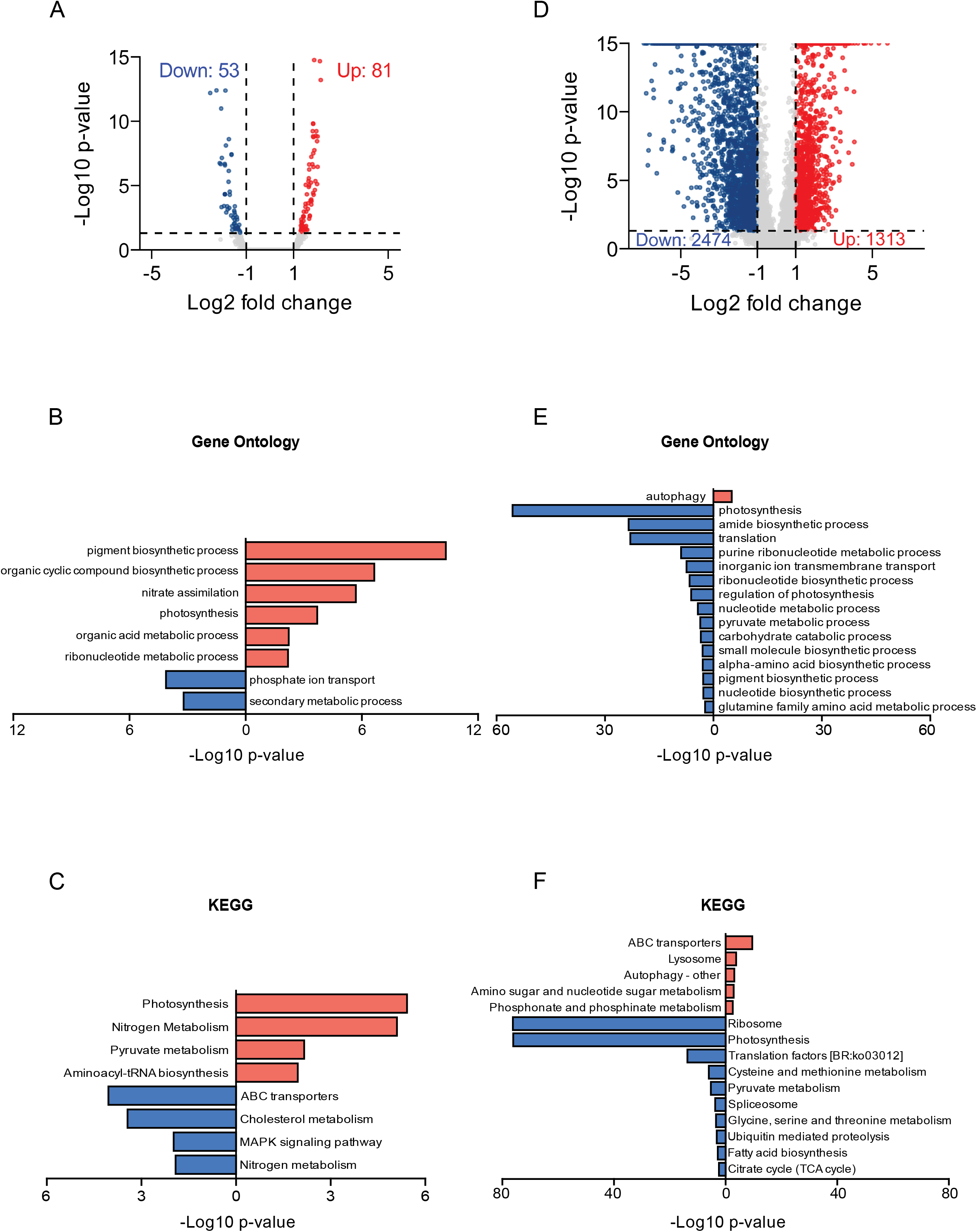
N-depletion of dinoflagellate *A. carterae* induces minor difference of mRNA accumulation level while dramatic change of translation efficiency. (A) Volcano plot showing differentially expressed genes (DEGs) under N-depletion treatment. (B) and (C), GO and KEGG enrichment analysis shows the enriched GO terms and pathways of these DEGs respectively. (D) Volcano plot showing differential translation efficiency genes (DTEGs) after N-depletion treatment. (E) and (F), Functional analysis of these DTEGs respectively. Red colors indicate either up-regulated DEGs or DTEGs, while blue colors represent either down-regulated DEGs or DTEGs.

**Figure S7.**
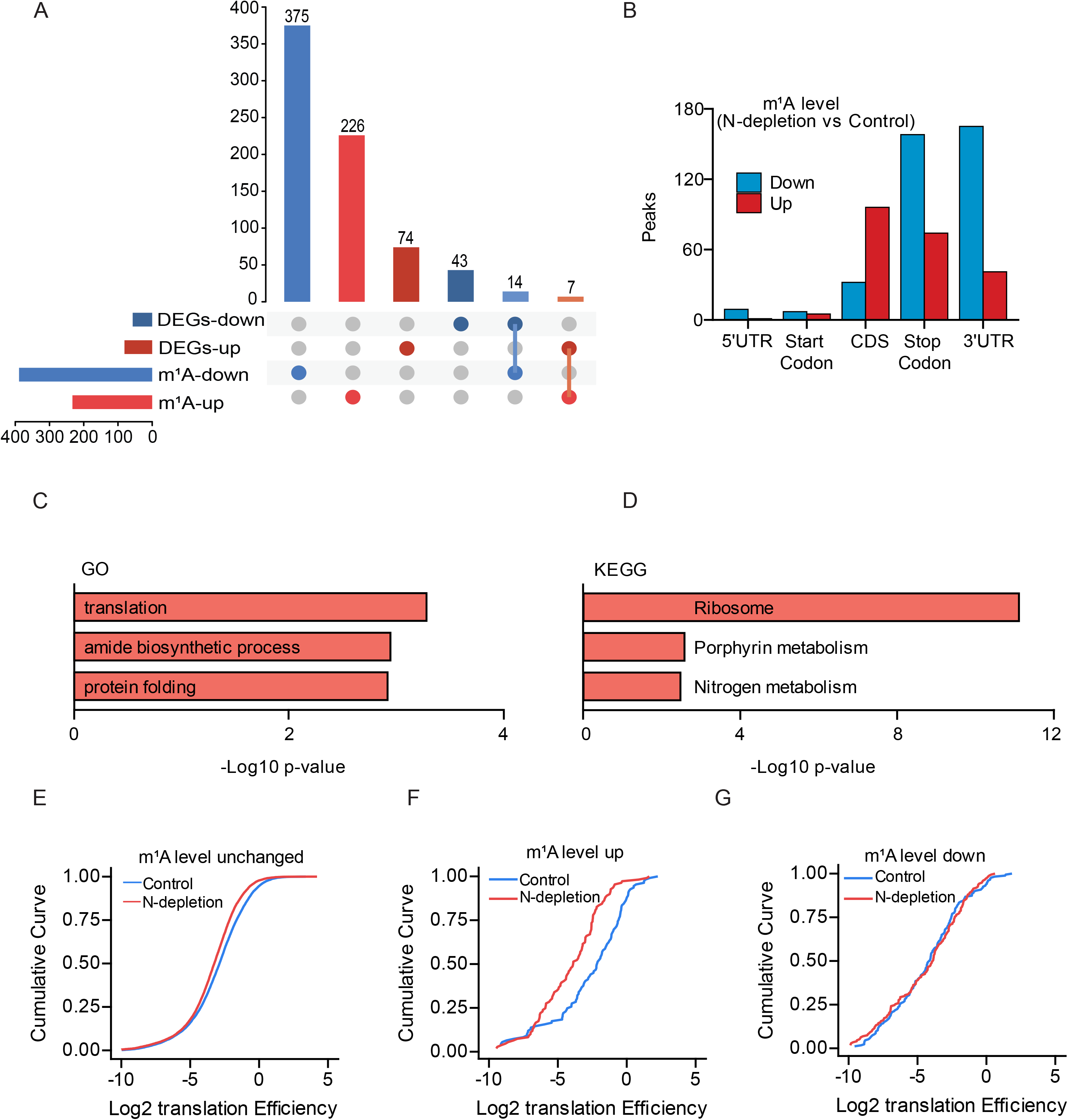
Features of differentially methylated m^1^A genes under N-depletion. (A) The Upset plot showing the numbers of shared genes between DEGs and differentially methylated genes. (B) The m^1^A peaks categories of differentially methylated genes along different mRNA segments. (C) and (D), GO and KEGG enrichment analysis of the upregulated m^1^A-modified genes. (E-G) The Log2 translation efficiency is plotted as accumulative fractions for transcripts with unchanged methylation (E), elevated methylation (F) and decreased methylation (G) for both control and N-depletion treatment groups.

**Figure S8.**
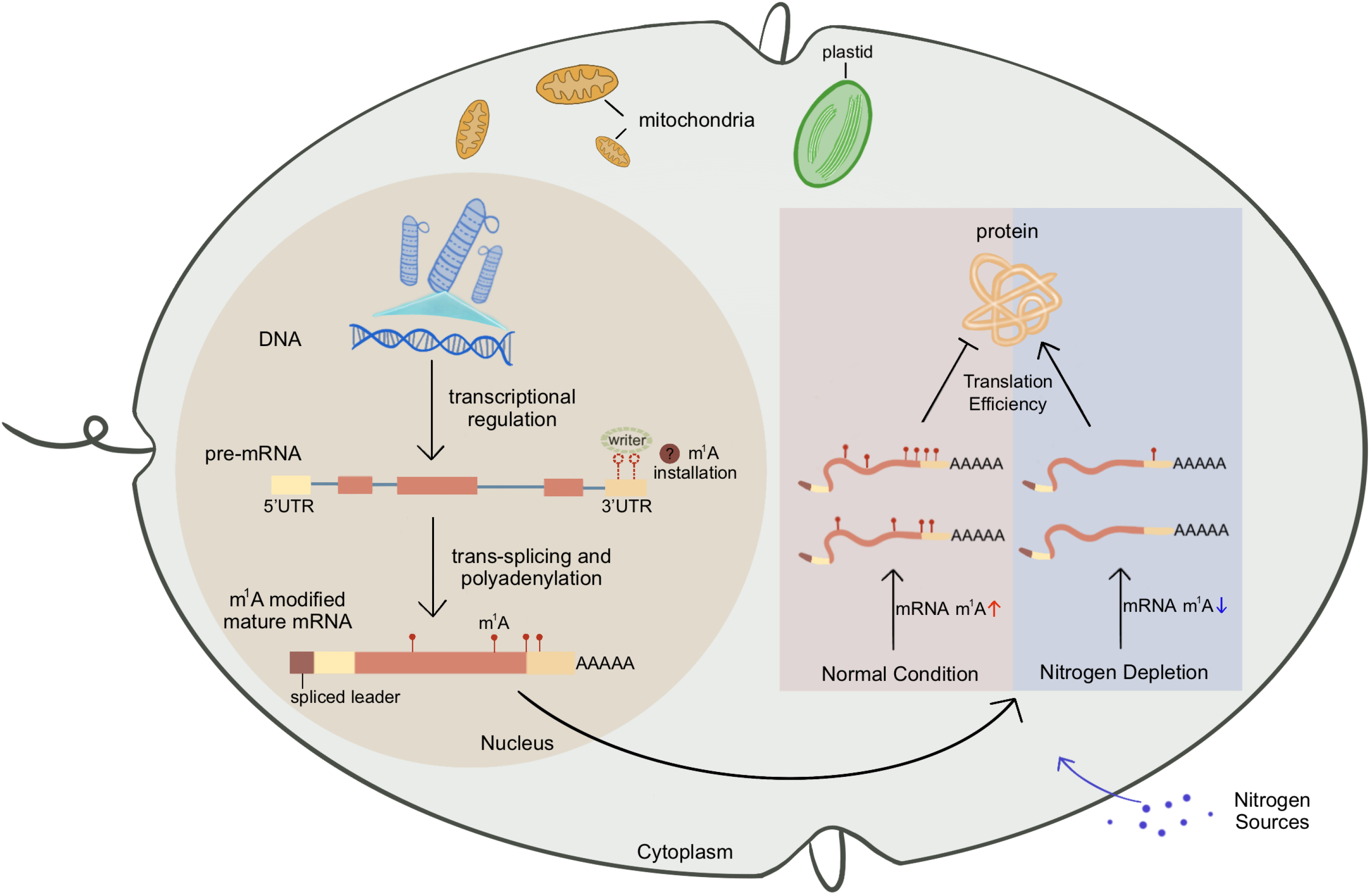
Working model for mRNA m^1^A modification as a critical layer of gene expression regulation in dinoflagellates.

## Materials and Methods

### Cell culture

Dinoflagellates *Amphidinium carterae* and *Symbiodinium* sp. were obtained from Shanghai Guangyu Biotechnology Co., Ltd. (Shanghai, China). The strains were normally cultured in f/2 medium at a constant temperature of 23°C. The lighting conditions were maintained under a 12:12 hour light-dark cycle with a photon flux density of 50µmol·m^-2^·s^-1^. Sub-culturing was performed at 7- to 10-day intervals to ensure optimal growth. The other dinoflagellate *Crypthecodinium cohnii* ATCC 30556 was purchased from American Type Culture Collection (ATCC) and routinely grown in A2E6 medium at dark conditions following official instruction. HEK293T cells were cultured in Dulbecco’s Modified Eagle’s Medium (DMEM, Gibco), supplemented with 10% fetal bovine serum (FBS, BI) and 1% penicillin-streptomycin (Thermo Fisher). The cells were maintained in a humidified incubator at 37°C with 5% CO_2_.

For nitrogen depletion experiment in *A. carterae*, the exponentially grown cells were washed once with sterile sea water and then cultured in normal and nitrogen-depletion f/2 medium respectively at an initial concentration of 2.5 × 10^4^ cells/ml. Cell numbers of *A. carterae* cells during the whole growth period were counted using automated Countess TM II FL cell counter (Thermo Fisher). *A. carterae* cells grown at exponential stage (day 6) was used to perform mRNA isolation and m^1^A content quantification.

### RNA preparation for LC-MS/MS analysis

Total RNA was extracted from *A. carterae*, *Symbiodinium sp.*, *C. cohnii* and HEK293T cell pellets using RNAzol®RT (MRC) according to the manufacturer’s protocols. The total RNA from *Chlamydomonas reinhardtii* received as a kind gift from Prof. Hongjie Shen, Fudan University. All materials and reagents used were RNase-free to minimize RNA degradation. Polyadenylated RNA (poly(A)+ RNA) was enriched from the total RNA samples by using two successive rounds of purification with VAHTS mRNA Capture Beads (Vazyme Biotech, Nanjing, China). RNA samples were digested by 50 units nuclease P1 (NEB) in a 25-µL reaction mixture containing nuclease P1 buffer at 37°C for 2 hours, followed by addition of 5 units Antarctic Phosphatase (NEB) and 2.5 µL 10 × Antarctic Phosphatase buffer. The enzymatic digestion was further conducted at 37°C overnight. The enzymatic reaction was terminated by adding 75µL of methanol. The samples were then centrifuged at 13000 rpm for 15 minutes. The supernatant was lyophilized using a vacuum freeze-dryer to concentrate the RNA digest and reconstituted in 50 µL of RNase-free water. 5 µL of samples was injected into LC–MS/MS system (Agilent6470 triple-quadrupole mass spectrometer or Q Exactive high-resolution benchtop quadrupole Orbitrap mass spectrometer) on an Agilent XDB-C18 column. Nucleosides were detected by using retention time and nucleoside to base ion mass transitions of m/z 268.1 to 136.1 (A), m/z 282.1 to 150.1, m/z 282.1 to 150.1 (m^6^A), m/z 285.1 to 153.1 (d3-m^1^A), m/z 298.1 to 166.1 (m^7^G) and m/z 296.1 to 150.1 (m^6^Am). Nucleoside standards with serial dilution (A, from 50 nM to 1600 nM; m^1^A, from 7.5 nM to 240 nM; m^6^A,2 nM to 64 nM) were run in parallel to produce the standard curves and obtain the target nucleosides concentration.

### Dot blot assay

The concentration of isolated total RNA and mRNA from both *A. carterae* and HEK293T cells were firstly determined using Equalbit RNA HS Assay Kit (Vazyme Biotech, Nanjing, China) and Qbit (ThermoFisher). An aliquot of each RNA sample (∼100 ng) was spotted onto a positively-charged nylon membrane and subsequently crosslinked with UV light using the chamber of SG linker. The membrane was then stained with methylene blue solution (0.2% methylene blue in 0.4 M sodium acetate and 0.4 M acetic acid) to ensure equal loading. After washing, the membrane was blocked with 5% milk in 1 × TBST at room temperature for 1-2 hours and incubated at 4°C overnight with a 1:1000 dilution of anti-m^1^A antibody (MBL #D345-3). On the second day, the membrane was washed with 1 × TBST for three times and then incubated in a 1:5000 dilution of goat anti-mouse IgG secondary antibody (Signalway Antibody LLC, Maryland, USA) for 1 hour at room temperature. After washing another three times in 1 × TBST, the blots were visualized with the Clarity Western ECL Substrate (BIO-RAD).

### m^1^A-RIP

200 ng poly(A)+ RNA of *A. carterae* was fragmented at 70°C for 10 min using 10 × RNA fragmentation reagent (Invitrogen) in a 10 µL reaction mixture. Subsequently, the fragmented mRNA was purified using RNA Clean & Concentrator-5 kit (Zymo Research). For m^1^A immunoprecipitation, each sample was processed as follows: Protein A/G Dynabeads (10 µL) were washed three times with 1 × IP buffer (150 mM NaCl, 0.1% NP-40, 10 mM Tris-HCl, pH 7.4) and then resuspended in 200 µL of 1 × IP buffer. Afterward, 1µL of anti-m^1^A antibody and 2.5 µL of protease inhibitor were added to the Protein A/G Dynabeads and incubated at 4°C for an hour. To enrich the m^1^A-modified mRNA, the antibody-coated beads were mixed with 200 µL of 1 × IP buffer, 2.5 µL of 1 × complete EDTA-free protease inhibitor (Roche Diagnostics), 1 µL of RNase inhibitor (Cwbio, Beijing, China) and fragmented mRNA from previous purification. 10 µL of the resultant mixture was retained as input sample. The mixture was incubated at 4°C with gentle rotation for 3 hours. Then, samples were washed sequentially with 1 mL of 1 × IP buffer, low-salt buffer (50 mM NaCl, 0.1% NP-40, 10 mM Tris-HCl, pH 7.4), and high-salt buffer (500 mM NaCl, 0.1% NP-40, 10 mM Tris-HCl, pH 7.4) at 4°C for 5 minutes. Elution of the immunoprecipitated products was performed with 5 units of Proteinase K in a buffer containing 5 mM Tris-HCl (pH 7.5), 1 mM EDTA, and 0.25% SDS at 37°C for 1.5 hours. Finally, the eluted RNA was purified with Zymo oligo clean and concentration kit and subjected to library preparation using the VAHTS Universal V8 RNA-seq Library Prep Kit for Illumina (Vazyme Biotech, Nanjing, China). Both the input and IP samples with three biological replicates independently were sent to Genewiz company (Jiangsu, China) for sequencing on an Illumina Novaseq 6000 platform with paired-end 150 bp mode.

### RNA-seq and Ribo-seq experiments

For RNA-seq, the isolated total RNA from *A. carterae* cells (three biological replicates for all treatments) was sent to the Beijing Genomics Institute (BGI, Shenzhen, China) for strand specific mRNA library construction and sequencing on DNBSEQ PE150 platform. On the other hand, to obtain the poly(A) tail length of *A. carteae* genes, one aliquot total RNA sample was sent to Wuhan Benagen Technology Co., Ltd. for nanopore direct RNA-sequencing using PromethION sequencer (Oxford Nanopore Technologies, Oxford, UK).

For Ribo-seq, *A. carterae cells* (∼40 million cells, two biological samples for each treatment) collected in the morning were treated with 100 µg/ml cycloheximide (CHX) for 8 minutes. After treatment, cells were centrifuged at 2000 rpm for 10 minutes at 4°C. The resulting cell pellets were washed once with CHX-containing 1 × PBS and resuspended in 2-3 volumes of lysis buffer composed of 10 mM Tris-HCl (pH 7.5), 100 mM KCl, 5 mM MgCl_2_, 1% Triton X-100, 100 µg/ml CHX, 2 mM DTT, and 1 × complete EDTA-free protease inhibitor (Roche Diagnostics), followed by sonication at a power of 100 W using an ultrasonic processor (SCIENTZ-950E, SCIENTZ, China) for 2 min (3s on / 3s off) in a ice-water bath. The supernatant was collected after centrifugation at 12000 g for 10 min at 4°C. The optical density (OD) of the cell lysate was determined from the absorbance of the supernatant at 260 nm with NanoDrop 2000. For Ribosome Protected Fragments (RPFs) generation, 15 U of RNase I (Invitrogen) was added per 1 OD of lysate, followed by a 40-minute incubation at 22°C with gentle mixing. RPFs were isolated using MicroSpin S-400 columns (Cytiva) and purified using the RNA Clean & Concentrator-25 kit (Zymo Research). After size-separation of RPFs using a 15% polyacrylamide TBE-urea gel (Coolaber Technology Co., Ltd, Beijing, China). Gel slices containing fragments of 17 to 34 nucleotides were excised using a scalpel. RNA recovery from the gel slices was performed using the ZR small-RNA PAGE Recovery Kit (Zymo Research). Purified RNA was mixed with 10 × T4 polynucleotide kinase reaction Buffer and T4 polynucleotide kinase and incubated at 37°C for 20 minutes with gentle mixing, followed by addition of 1 × T4 DNA ligase buffer and another incubation at 37°C for 20 minutes. The samples were repurified using the RNA Clean & Concentrator-25 kit (Zymo Research) and proceeded directly to cDNA library construction using the VAHTSTM Small RNA Library Prep Kit for Illumina (Vazyme Biotech, Nanjing, China). Sequencing was performed on the BGI MGISEQ-2000 platform using PE50 strategy.

### Isolation of poly(A) tail and non-poly(A) segments from *A. carterae* poly(A)+ RNA

200ng of purified poly(A)+ RNA from *A. carterae* was first fragmented at 70°C for 10 min using 10 × RNA fragmentation reagent (Invitrogen) in a 10 µL reaction mixture. Following fragmentation, the poly-A tails were captured using VAHTS mRNA Capture Beads (Vazyme Biotech, Nanjing, China). Meanwhile, the non-poly(A) RNA segments in the flow-through were purified using the RNA Clean & Concentrator-5 kit (Zymo Research). The resulting purified poly(A) tails and non-poly(A) segments were then analyzed using an anti-m^1^A antibody-based dot-blot assay.

### Protein purification

The encoding sequences of both AcTRMT6 and AcTRMT61 were optimized with *Escherichia coli* codons, synthesized by Tsingke Biological Technology Co., Ltd. (Beijing, China) and inserted into the expression vector pETDuet-1 to generate pETDuet-1-AcTRMT-AcTRMT61 plasmid, which was then transformed into *E. coli* BL21 (DE3) competent cells. Cells were cultured at 37°C until OD600 reaches 0.6 - 0.8, then induced with 0.1 mM IPTG and incubated at 16°C for 16 hours. The harvested cells were lysed using high-pressure homogenizer (AH-NANO, ATS, China) with a pressure of 800 ∼ 900Lbar for four cycles in the lysis buffer (20 mM Tris–HCl, pH 7.4, 300 mM NaCl, 15mM imidazole, 0.25% TritonX-100, 10% glycerol). The lysate was centrifuged at 20,000 rpm for 30 minutes, and the supernatant containing 6× His-tagged AcTRMT-AcTRMT61 complex was purified using Ni-NTA beads (Changzhou Smart-Lifesciences Biotechnology Co., Ltd., China) following the manufacturer’s guidelines. In brief, the protein-bound beads were washed three times with wash buffer (20 mM Tris–HCl, pH 7.4, 300 mM NaCl, 25 mM imidazole, 10% glycerol) and then eluted using the elution buffer (50 mM Tris-HCl, pH 7.4, 150 mM NaCl, 250 mM imidazole and 10% glycerol). Eluted proteins were pooled and dialyzed extensively with dialysis buffer (20 mM Tris–HCl pH 7.4, 50 mM NaCl, 0.5 mM EDTA, 1 mM DTT, and 10% glycerol). All purification steps were conducted at 4°C. The purity and size of the isolated protein complex were confirmed through Coomassie blue staining of sodium dodecyl sulfate-polyacrylamide gels.

### Biochemical assay of AcTRMT6-AcTRMT61 heterocomplex activity *in vitro*

Biochemical assays of the purified AcTRMT6-AcTRMT61 enzymes were conducted in 50 µL reaction mixtures at 28°C overnight. Each mixture contained 1× MTase buffer (100 mM Tris, pH 8.0, 1.0 mM DTT, 0.1 mM EDTA, 10 mM MgCl_2_, 20 mM NH_4_Cl), 32 µM d3-SAM (deuterium-labeled S-Adenosylmethionine), 5 µM recombinant protein complexes, and various substrates. Tested substrates included total RNA, mRNA, and small RNA from *A. carterae*, as well as total RNA and mRNA from HEK293T cells. The small RNA (< 200 nt) from *A. carterae* was purified using the RNA Clean & Concentrator-25 kit (Zymo Research). For control, enzyme-inactivation reactions through heat treatment were set up using total RNA from *A. carterae* cells. After overnight incubation, the reaction mixtures were heat-inactivated at 95°C for 5 minutes and cooled down in an ice bath for another 5 minutes. Nuclease P1 (NEB) and Antarctic Phosphatase (NEB) were then added to digest the samples for subsequent LC-MS/MS analysis.

### Bioinformatics analyses of RNA-seq, Ribo-seq and MeRIP-seq data

A *de novo* genome assembly of *A. carterae* (Acart.genome.v1, unpublished dataset) based on HiFi reads of Pacific Biosciences Sequel II system was used as the reference genome in this study. Raw reads data of RNA-seq, Ribo-seq and methylated RNA immunoprecipitation sequencing (MeRIP-seq) were prepared according to the procedure describe previously (Chothani, Adami et al., 2022). Briefly, TrimGalore (version 0.6.7) was used to remove adapters and low-quality base (-q 25) at first. Of note, for Ribo-seq down-stream analysis, reads with length in the range of 20-35 bp were kept. RPFs in Ribo-seq represents actively translated mRNA transcript and thus reads mapping to rRNA sequences were discarded. The trinucleotide periodicity of ribosomes and codon usage frequency were examined using revised RiboseQC package (v0.99) (Calviello, Sydow et al., 2019).

Pre-processed reads of RNA-seq, Ribo-seq and MeRIP-seq were then subjected to alignment against Acart.genome.v1 using software STAR (version 2.7.10a) (Dobin, Davis et al., 2012), allowing for no more than 2 mismatch. The sorted mapping files of Ribo-seq and RNA-seq were used for quantification of with StringTie (v2.1.7) (Pertea, Pertea et al., 2015). Note that Ribo-Seq reads were mapped with additional parameters: --outFilterScoreMinOverLread 0.33, --outFilterMatchNminOverLread 0.33. Next, the obtained read counts were converted to TPM (transcripts per million). Genes with cutoff values of TPM > 5 for RNA-Seq were used for downstream analysis. The R package DESeq2 was used to perform differential gene expression of RNA-seq datasets. Next, translational efficiency (TE) analysis was calculated by dividing the ribosome footprint density using a Ribo-seq TPM value larger than 0 with mRNA abundance (RNAseq TPM) (Chothani et al., 2022). The differential translation efficiency gene (DTEG) was analyzed by RiboDiff (v0.2.2)(Zhong, Karaletsos et al., 2016). After alignment, m^1^A peaks enriched via IP over input control were detected based on a combination of corresponding biological replicates using MACS2 (v2.2.7.1)(Zhang, Liu et al., 2008) algorithm with -q 0.05. Minimum free energy (MFE) calculation of m^1^A methylated genes was performed by using the ViennaRNA package (Lorenz, Bernhart et al., 2011).

Motifs within these m^1^A peaks were determined by HOMER software (Heinz, Benner et al., 2010). Differential methylation sites with a fold change of > 1.5 and p value of < 0.05 were identified using DiffBind (Stark & Brown, 2012) with edgeR (Robinson, McCarthy et al., 2010). All GO (Gene Ontology) terms and KEGG (Kyoto Encyclopedia of Genes and Genomes) pathway enrichment analyses were performed by R package clusterProfiler (v4.2.2)(Yu, Wang et al., 2012)

### Definition of m^1^A peaks in different mRNA segments

The locations of m^1^A peaks were classified into five categories based on their overlap with mRNA segments using BEDTools (Quinlan & Hall, 2010). Specifically, when a peak overlaps with either the start codon or strop codon, we assigned these peaks into “start codon” and “stop codon” regions respectively. Otherwise, these peaks were considered to localize in 5L UTR, CDS and 3L UTR respectively according to their positions.

